# In Humans, fMRI Reveals That Striosome-like and Matrix-like Striatal Voxels are Engaged in Different Phases of Movement

**DOI:** 10.1101/2025.09.01.673509

**Authors:** Alishba Sadiq, Jeff L. Waugh

## Abstract

**Introduction:** The striatum is organized into two neurochemically and anatomically distinct compartments, the striosome and matrix, that play specialized roles in motor and cognitive functions. While extensive animal research has elucidated compartment-specific contributions to reward, learning and motor control, direct evidence for compartment specialization in humans is lacking.

**Methods:** We defined human striatal voxels as striosome-like or matrix-like based on biases in structural (diffusion) connectivity. Then we investigated functional activation patterns in those compartment-like voxels using task-based functional MRI (tfMRI) during pre-movement cue and five motor conditions (left/right hand, left/right foot, and tongue movements).

**Results:** Functional activation was strikingly segregated: striosome-like voxels were preferentially engaged during the cue phase, while matrix-like voxels dominated activation during motor execution, especially for tongue and foot movement. Motor tasks elicited robust bilateral activation, with contralateral activation dominating during limb movements. Activation was more lateralized in matrix-like than in striosome-like voxels. Both striosome-like and matrix-like voxels exhibited strong activation at the onset of task execution (e.g., within the first few seconds post-cue). However, activation in matrix-like voxels declined modestly over the course of the movement phase, while striosomal activation dropped sharply at task termination, suggesting a role in behavioral transitions. These findings are consistent with the role of the striosome in anticipatory evaluation and dopaminergic modulation, and matrix specialization for executing automatized routines.

**Conclusions:** This study provides the first task-based fMRI evidence of temporally and functionally distinct striatal compartment dynamics in humans, offering novel insights into striatal microcircuitry in motivated behavior and the planning and execution of movements.

**Key Points:**

- Striatal medium spiny neurons develop in two interdigitated tissue compartments, the striosome and matrix, that are embryologically, pharmacologically, and anatomically distinct. Inter-compartmental differences in function have been identified in animals, but never in humans.
- We found that in humans, the compartments differed in functional activation during movement tasks: during the task cue, activation was greater in striosome-like voxels, while matrix-like activation was greater during each of five distinct types of movement.
- Both compartments were active at the beginning of movement, but at the termination of movement striosome-like activation fell to below baseline, suggesting a role for the striosome in behavioral transitions.

## 1 Introduction

The basal ganglia are a subcortical cluster of interconnected nuclei, historically implicated in motor control, but now recognized to contribute to skill acquisition and action selection in executive, emotional, learning, and reward tasks (Bamford and Bamford 2019; Rocha, Freire et al. 2023). While dysfunction of these circuits was classically associated with movement disorders such as Parkinson disease, Huntington disease, dystonia, and dyskinesias (Wichmann and Dostrovsky 2011), striatal pathology has also been implicated in a wide range of neuropsychiatric and cognitive disorders, including obsessive-compulsive disorder (OCD; Burguiere, Monteiro et al. 2015), Tourette syndrome (TS; Albin 2006), addiction, and depression (Dunlop and Nemeroff 2007; Kalivas 2008).

The striatum is the principal input nucleus of the basal ganglia and is the recipient of cortical, thalamic, and brainstem efferents, each of which contributes to the selection and execution of discrete tasks. Striatal projection neurons (SPNs) can be subdivided into two tissue compartments, striosome and matrix, that have distinct embryologic origins, histochemistry, pharmacology, and both afferent and efferent connectivity (Crittenden and Graybiel 2011). The striosome receives input from the limbic cortices and directly inhibits the dopaminergic neurons of the substantia nigra pars compacta, while both striosome and matrix compartments receive nigral dopaminergic inputs (Watabe-Uchida, Zhu et al. 2012; Friedman, Homma et al. 2015; Crittenden, Tillberg et al. 2016). This bidirectional connectivity provides a basis for their differential dopamine-related modulation: dopaminergic inputs decrease striosome firing but increase matrix firing (Prager, Dorman et al. 2020). Such inverted dopaminergic responses, combined with the distinct afferent and efferent connectivity of the compartments, suggest that the striosome may dynamically regulate motivational state and reward-based evaluation through direct modulation of midbrain dopamine neurons, whereas the matrix may facilitate the initiation and execution of appropriate motor actions via its connections to sensorimotor pathways. Matrix SPNs receive input from sensorimotor and associative cortices and project predominantly to the globus pallidus interna and substantia nigra pars reticulata (Lévesque and Parent 2005; Watabe-Uchida, Zhu et al. 2012), highlighting the role of the matrix in sensorimotor integration and motor control. Striosome and matrix compartments appear to be functionally distinct yet complementary, as they are differentially engaged during processes such as value-driven learning, affective evaluation, and action selection (Weglage, Wärnberg et al. 2021). Recent evidence suggests that both compartments participate in motor control but they do so via separate and potentially opposing circuits (Okunomiya, Watanabe et al. 2025). Specifically, striosomal direct pathway SPNs (dSPNs) suppress movement and dopamine release at the end of a movement epoch, whereas matrix dSPNs promote movement, underscoring the complementary yet distinct contributions of the two compartments to the regulation of movement.

While this compartmental architecture has been well-described in animal models and human histology, its functional significance during active motor behavior remains poorly understood. Selective ablation of the striosome impairs motor skill learning while sparing general motor functions (Lawhorn, Smith and Brown 2009). Conversely, matrix neurons have been directly implicated in the execution of motor behavior. Movement-related SPNs in rodents localize predominantly within the matrix or along matrix– striosome borders, reinforcing the matrix’s central role in motor execution (Trytek, White et al. 1996). Consistent with this, selective disruption of GABA release from matrix SPNs produced pronounced motor abnormalities, including hyperactivity and muscle rigidity, as well as non-motor effects such as growth retardation and memory deficits. These phenotypes worsened with age, underscoring the critical role of intact GABAergic transmission in matrix SPNs for maintaining motor control and behavioral stability (Reinius, Blunder et al. 2015).

Further evidence of compartment-specific functions comes from Huntington disease models. In the YAC128 mouse model (Lawhorn, Smith and Brown 2008), loss of volume and SPN number was greater within the striosome, relative to the matrix, and this degeneration strongly correlated with deficits in motor coordination and balance. This finding does not contradict the essential role of the matrix in motor execution; rather, it suggests that striosome degeneration can secondarily impair motor control, likely through its modulatory influence on dopaminergic signaling and striatal circuit balance, thereby contributing indirectly to motor dysfunction. While the matrix is primarily implicated in the direct execution of movement, these findings suggest that the striosome plays a critical modulatory role in fine motor tuning, motor learning, and the integration of affective or contextual signals. This view aligns with recent findings that the two compartments regulate *different phases of movement*: striosomal dSPNs dampen activity at the end of a movement epoch by suppressing dopamine release, while matrix dSPNs sustain motor execution. These results highlight not only opposing roles, but also complementary temporal dynamics, with striosome dysfunction likely disrupting the smooth transition between motor states (Okunomiya, Watanabe et al. 2025). Beyond neurodegenerative models, functional imbalances between the compartments have been linked to abnormal behavior. Canales & Graybiel (2000) demonstrated that in rats, the ratio of striosome:matrix activation accurately predicted the severity of habitual motor movements following doses of cocaine or methamphetamine, suggesting that imbalances in the relative function of striosome and matrix can drive repetitive motor patterns and behavioral inflexibility (Canales and Graybiel 2000). Collectively, these findings indicate not only that compartmental imbalances can produce motor impairments, but also that striosome and matrix may be dissociably involved in different facets of motor control: the matrix facilitating efficient motor output, and the striosome modulating the motivational, contextual, or evaluative dimensions of movement. These findings suggest that imbalances between striosome and matrix activity contribute to motor impairments, reinforcing the importance of studying the independent role of each compartment in the regulation of movement.

Decades of prior animal studies suggested a functional division of labor between the compartments. The striosome is thought to regulate anticipatory and evaluative processes such as action selection, informed by reward potential or limbic state, whereas the matrix is considered critical for executing well-learned, habitual motor actions (Graybiel 2008; Hikosaka, Kim et al. 2014; Friedman, Homma et al. 2015). While this framework was formulated based on structural connectivity patterns (Donoghue and Herkenham 1986; Flaherty and Graybiel 1993; Eblen and Graybiel 1995), subsequent functional studies provided direct behavioral evidence for striosome involvement in reward evaluation and decision-making. For example, placement of stimulating electrodes in the striosome, but not in the matrix, led to habitual self-stimulation (White and Hiroi 1998). Similarly, striosomal activation has been linked to pessimistic decision-making under emotional conflict (Amemori, Graybiel and Amemori 2021) and suppression of movement under threat (Friedman, Homma et al. 2015), highlighting the compartment’s role in affectively guided behavioral control.

We recently demonstrated that connectivity-based parcellation (probabilistic diffusion tractography) can identify voxels with striosome-like and matrix-like properties (eg, biases in structural connectivity, intrastriate location) in living humans (Waugh, Hassan et al. 2022). This method enables, for the first time, individualized mapping of compartment-like voxels *in vivo*, based on their distinct cortical and subcortical connectivity profiles. We refer to the parcellated striatal voxels as “striosome-like” or “matrix-like” to emphasize the inferential basis of this approach and to clarify that these voxels are not direct equivalents of the striosome and matrix compartments identified by immunohistochemical analyses. However, our MRI-based parcellations closely mirror key anatomical features of the striosome and matrix compartments observed in tissue. First, the relative abundance of matrix-like and striosome-like voxels matched histological estimates, with matrix-like voxels comprising approximately 85% of the striatum and striosome-like voxels approximately 15%, a ratio consistent with prior human and primate tissue studies (Desban, Kemel et al. 1993; Holt, Graybiel and Saper 1997; Waugh, Hassan et al. 2022; Funk, Hassan et al. 2023; Funk, Hassan and Waugh 2024; Sadiq, Funk and Waugh 2025). Second, the spatial distribution of compartment-like voxels conformed to the distribution of the compartments in tissue, with striosome-like voxels preferentially localized to the rostroventral striatum, and matrix-like voxels more evenly distributed, especially in the dorsolateral and caudal striatum (Graybiel and Ragsdale Jr 1978). Third, projections to striosome-like and matrix-like voxels are organized somatotopically (Funk, Hassan et al. 2023), just as they are in animals (Eblen and Graybiel 1995). Fourth, matrix-like voxels are more likely to occur in large, contiguous clusters, while striosome-like voxels are more likely to occur in isolated clusters. Finally, connectivity-based parcellations are highly reproducible between repeated MRI scans, with a 0.14% test-retest error rate (Waugh, Hassan et al. 2022). These connectivity-based parcellations lay the foundation for investigating human striatal compartmental function in both rest and task-based fMRI experiments (Sadiq, Funk and Waugh 2025).

However, the extent to which these compartment-specific functions translate to human behaviors remains largely unknown. We previously demonstrated that striosome-like and matrix-like voxels participate in distinct structural networks (Funk, Hassan et al. 2023; Funk, Hassan and Waugh 2024) and are embedded in segregated resting-state fMRI networks (Sadiq, Funk and Waugh 2025). This compartmental segregation aligns with prior hypotheses that striosomes and matrix may be differentially implicated in neuropsychiatric and movement disorders. In particular, (Crittenden and Graybiel 2011) reviewed evidence suggesting that disorders such as Huntington disease, Parkinson disease, depression, and OCD may exhibit selective vulnerability or dysregulation of the striosome system, potentially due to its unique connectivity with limbic networks and nigral dopaminergic neurons. However, direct evidence of functional dissociation during active behavior in humans has not been demonstrated previously, to the best of our knowledge. To address this gap, we leveraged our connectivity-based parcellation method to examine functional activation in striosome-like and matrix-like voxels during simple motor tasks. Using these compartment-specific masks as seeds in a task-based fMRI design, we tested whether striosome and matrix compartments show distinguishable activation profiles during motor behaviors. We aimed to answer three critical questions: (1) Do striosome and matrix compartments exhibit functionally distinct activation patterns in the human brain during motor tasks? (2) Are these differences specific to phases of the task, e.g. during cue-related or motor execution periods? (3) Do these dynamics support the proposed roles of the striosome in anticipatory processing and of the matrix in habitual motor control?

By comparing activation in compartment-like voxels during the cue, initiation, plateau, and termination phases of movement, we provide the first evidence that striosome and matrix differ in both the amplitude and timing of task-evoked functional responses in humans. These findings have important implications for understanding striatal microcircuit contributions to motor planning, action execution, and the pathophysiology of conditions in which dysfunctional striatal signaling has been implicated, such as OCD, dystonia, and tic disorder.

This study provides the first direct evidence of functional dissociation between striosome- and matrix-like activation in the human striatum during motor behaviors, offering a new framework for linking microcircuit-specific dynamics to complex motor and neuropsychiatric symptoms.

## 2 Materials and Methods

Overview: we used a combination of structural and functional connectivity to investigate compartment-specific activation during motor tasks in humans (Figure 1). We first utilized connectivity-based parcellation (probabilistic diffusion tractography) to identify striatal voxels with striosome-like or matrix-like biases in structural connectivity. Notably, we have previously described striatal parcellation in 14 distinct human neuroimaging datasets (Waugh, Hassan et al. 2022; Funk, Hassan et al. 2023; Funk, Hassan and Waugh 2024; Sadiq, Funk and Waugh 2025) – in each dataset, compartment-like voxels followed the spatial distribution, relative abundance (striosome:matrix volume ratio), and extra-striatal connectivity patterns of striosome and matrix tissue identified through prior animal and human histological studies. Next, we examined functional activation within these compartment-like striatal masks, modeling each subject’s BOLD response to task events using a general linear model (GLM), a standard statistical approach that estimates brain activity related to specific task phases (e.g., movement initiation or execution). For each voxel, we computed contrast parameter estimates (COPEs) that quantified the difference in activation between task and baseline conditions, yielding compartment-specific activation metrics. This approach allowed us to directly compare the amplitude and timing of activation within striosome- and matrix-like voxels across different motor tasks.

**FIGURE 1:**
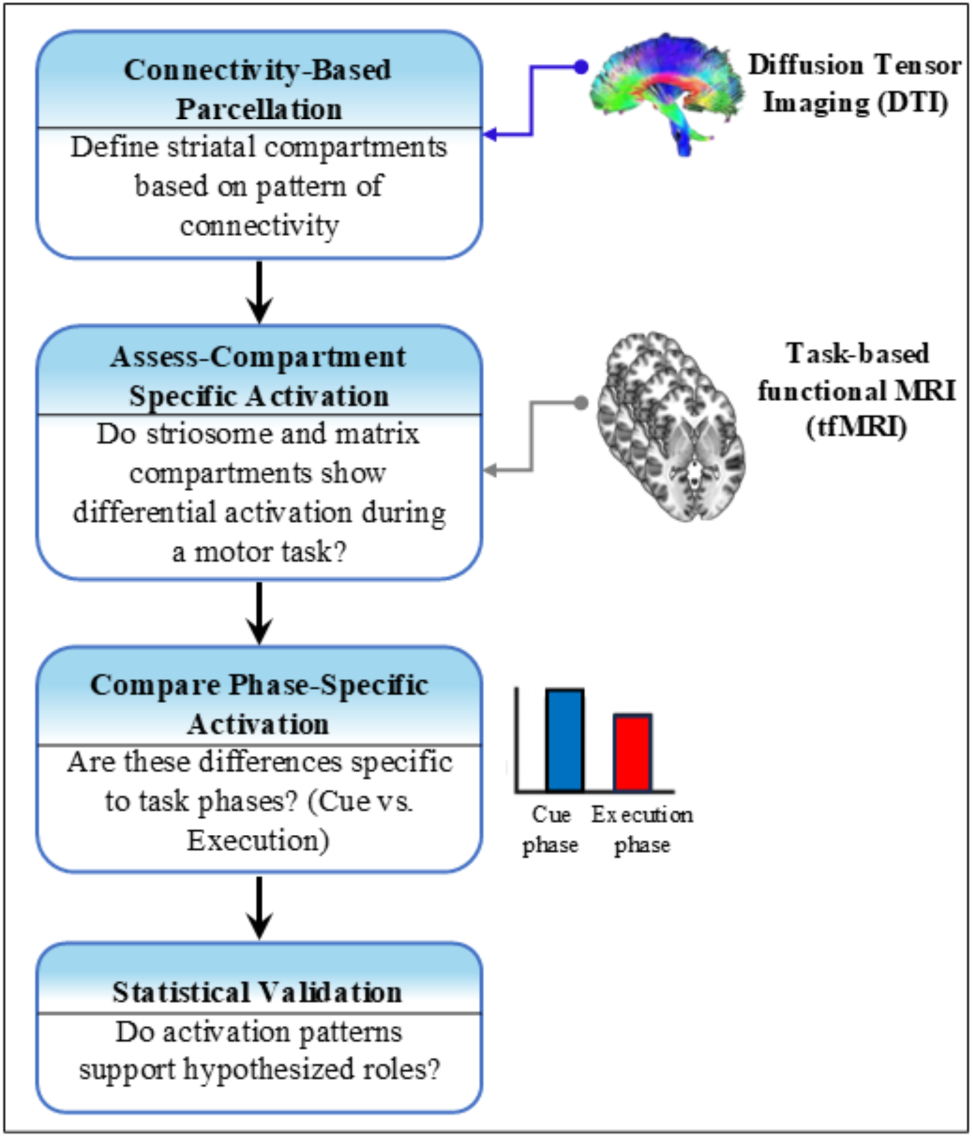
Overview of the experimental workflow for assessing compartment- and phase-specific activation in the human striatum. We used structural connectivity-based parcellation (probabilistic tractography based on diffusion tensor imaging [DTI]) to define striosome-like and matrix-like compartments, for each individual. Note that the precise location of the striosome varies between individuals – a uniform region-of-interest approach will not accurately distinguish the striatal compartments. These compartment masks served as regions of interest for evaluating compartment-specific activation using task-based functional MRI (fMRI) during motor performance. We compared cue-related and execution-related activity across compartments to test hypothesized roles of the striosome in anticipatory processing and the matrix in motor execution. Activation values are presented in arbitrary units (AU), reflecting relative BOLD signal change.

### 2.1 Study Population

This was a secondary analysis of MRI data from the Human Connectome Project (HCP) S1200 release (Van Essen, Smith et al. 2013), which included comprehensive behavioral and neuroimaging datasets from a large cohort of healthy young adults. From the original sample of 1,206 participants, we selected individuals who had complete diffusion MRI and task-based (motor) fMRI datasets. We excluded participants if they had any lifetime history of illicit or addictive substance use (cocaine, hallucinogens, cannabis, nicotine, opiates, sedatives, or stimulants) based on HCP diagnostic screening. Additionally, we excluded individuals who met the DSM-5 criteria for Alcohol Use Disorder (either Alcohol Abuse or Dependence) or who reported consuming more than four alcoholic drinks per week on average during the year prior to scanning. After these exclusions, our final study cohort comprised 701 healthy adults (mean age = 29.3 years, SD = 3.8), including 409 females and 292 males. All participants provided written informed consent at enrollment with the HCP study (Van Essen, Ugurbil et al. 2012). In a recent study whose experimental cohort overlapped with the cohort utilized here, we identified the resting state functional-networks that covaried with striosome-like vs matrix-like voxels (Sadiq, Funk and Waugh 2025).

### 2.2 MRI Acquisition Protocols

Task-based functional MRI (tfMRI) and diffusion tensor imaging (DTI) data were acquired as part of the Human Connectome Project (HCP) S1200 release using 3T scanners with harmonized imaging protocols across multiple sites. The tfMRI sessions used the same echo-planar imaging (EPI) parameters as resting-state fMRI (rfMRI), ensuring consistency in spatial and temporal resolution. Each tfMRI scan was acquired with a multiband gradient-echo EPI sequence with the following parameters: TR = 720 ms, TE = 33.1 ms, flip angle = 52*^◦^*, FOV = 208 *×* 180 mm, matrix = 104 *×* 90, with 72 axial slices and a multiband factor of 8. Echo spacing was 0.58 ms, and the bandwidth was 2290 Hz/pixel. The tfMRI resolution was 2.0 mm isotropic, which we have previously demonstrated is sufficient to resolve striosome-like and matrix-like structural connectivity (Waugh, Hassan et al. 2022). Each motor task condition included two runs of 284 frames, each lasting approximately 3 minutes and 34 seconds. The short scan duration and fast TR enabled high temporal resolution for task-evoked activity mapping. DTI data for S1200 subjects was acquired at 1.25 mm isotropic resolution using 200 directions (14 B0 volumes, 186 volumes at noncolinear directions) with the following parameters: repetition time = 3.23 s, echo time = 0.0892. DTI scans included both anterior-posterior and posterior-anterior acquisitions, allowing for correction of susceptibility artifacts.

#### 2.2.1 Motor-Task Design

The motor task paradigm was adapted from protocols developed by Buckner and colleagues (Buckner, Krienen et al. 2011; Yeo, Krienen et al. 2011) to robustly activate motor and somatosensory cortical and subcortical areas. Participants were visually cued to perform a series of specific movements, isolated to one body part at a time. Specifically, in separate blocks, participants tapped the left fingers, right fingers, moved the left toes, right toes, and tongue. Each block included a 3-second cue phase to prepare for the movement task, then 12 seconds of performing one of 10 discrete movements. Each run included 10 movement blocks: two tongue blocks, four hand blocks (two left, two right), and four-foot blocks (two left, two right). Within each run, three 15-second fixation blocks served as a baseline condition, during which participants fixated on a centrally presented cross without performing any movement. All motor task events were modeled relative to the average of the three 15-second fixation blocks, which served as a baseline condition. During these fixation periods, participants fixated on a centrally presented cross without performing any movement. This approach allowed us to contrast movement-related activation with a stable, aggregated baseline estimate.

### 2.3 Striatal Parcellation

We previously established a technique to identify striosome-like and matrix-like voxels in the human striatum based on their distinct *in vivo* connectivity profiles (Waugh, Hassan et al. 2022). We use the terms “striosome-like” and “matrix-like” to remind readers that these parcellations are inferential and are not the equivalent of immunohistochemical staining, the gold standard for identifying striosome and matrix in tissue. We constructed composite target masks of regions whose striatal structural connectivity was biased toward one compartment, as demonstrated through prior studies that utilized injected tract tracers in animals, or regions whose structural connectivity biases were demonstrated in human diffusion tractography (Waugh, Hassan et al. 2022). Striosome-favoring regions included the posterior orbitofrontal cortex, anterior insula, basolateral amygdala, basal operculum, and posterior temporal fusiform cortex. Matrix-favoring regions included the inferior frontal gyrus pars opercularis, primary motor cortex, supplementary motor area, primary somatosensory cortex, and superior parietal cortex.

We conducted striatal parcellation using the FSL tool *probtrackx2*, evaluating the relative connectivity of each striatal voxel to striosome-favoring vs. matrix-favoring target masks. We performed tractography in each subject’s native diffusion space and utilized standard parameters: curvature threshold = 0.2; steplength = 0.5 mm; number of steps per sample = 2,000; number of samples per seed voxel = 5,000; distance correction, to prevent target proximity from influencing connection strength. For each striatal voxel, we compared the number of streamlines reaching striosome-favoring versus matrix-favoring target regions. The resulting ratio of these seed-to-target streamline counts served as an index of compartmental bias. Each striatal voxel therefore had a bias probability of P=0-1. We defined compartment-specific connectivity bias as P>0.55 toward striosome-favoring or matrix-favoring target masks. This voxelwise comparison yielded a continuous map of striosome-like and matrix-like connectivity, specific to each subject and hemisphere. Notably, since the striosome is uniquely located in each individual, striosome-like and matrix-like masks must be uniquely located for each participant; standardized striatal region-of-interest masks are inadequate for investigating the striatal compartments.

Since each diffusion voxel has the potential to include both striosome and matrix, many striatal voxels have only modest compartment-like connectivity bias. To maximize the contrast between striosome-like and matrix-like voxels, we excluded voxels with indeterminate or low bias by applying an iterative thresholding approach to the compartment-like probability maps. For each subject and hemisphere, we selected voxels from the most-biased end of the distribution, representing either striosome-like or matrix-like connectivity, and gradually reduced the threshold until the total mask volume matched our target volume. We set this target at 13% of the original striatal mask, corresponding to 1.5 standard deviations above the mean in a Gaussian distribution. To ensure comparability between the compartments, we assured that striosome-like and matrix-like masks for each subject and hemisphere had equal volume. These high-bias, equal-volume masks have previously been shown to recapitulate the known spatial distribution and connectivity of striosome and matrix compartments, as established in histological studies (Waugh, Hassan et al. 2022; Funk, Hassan et al. 2023; Funk, Hassan and Waugh 2024; Sadiq, Funk and Waugh 2025). Then we registered each high-bias compartment-like mask from diffusion space to the subject’s structural (T1-weighted) image using FSL’s FLIRT, and then non-linearly transformed into fMRI space using the precomputed functional-to-structural registration matrices from the HCP pipeline. All transformations were visually inspected for accuracy. The resulting striosome-like and matrix-like masks in fMRI space were used for subsequent functional connectivity analyses.

### 2.4 Validation of Striatal Parcellation

To evaluate whether our parcellated voxels reflected known histological compartmentalization patterns, we measured the Cartesian position of every voxel within the equal-volume 1.5SD masks. For each subject and hemisphere, the Cartesian coordinates (x, y, z) of each voxel were extracted and referenced to the centroid of the corresponding nucleus (caudate or putamen). Spatial distribution was quantified by calculating within-plane dispersion and root-mean-square (RMS) distance from the nucleus centroid, providing a direct voxelwise measure of how striosome-like and matrix-like voxels were organized within three-dimensional striatal space. Notably, we have previously demonstrated that striosome-like voxels were consistently enriched in rostral, medial, and ventral regions of the striatum, in line with prior histological characterizations (Graybiel and Ragsdale Jr 1978; Goldman-Rakic 1982; Donoghue and Herkenham 1986; Ragsdale Jr and Graybiel 1990; Desban, Kemel et al. 1993; Eblen and Graybiel 1995; Waugh, Hassan et al. 2022).

Striosomal branches are embedded within the surrounding matrix (Graybiel and Ragsdale Jr 1978; Holt, Graybiel and Saper 1997). In coronal tissue sections, striosome branches appear as discrete “islands” amid a contiguous “sea” of matrix tissue. To quantify the spatial distribution of striosome-like and matrix-like voxels, we applied the fsl-cluster command, using a higher bias threshold of *P* > 0.87 to isolate voxels with high compartment-specific bias. We focused on the largest cluster within each compartment, which represented the dominant spatial organization of striosome- and matrix-like patterns.

We previously found that minor shifts in voxel location were enough to disrupt compartment-like structural connectivity patterns, indicating that such biases were dependent on precise voxel locations rather than their ‘neighborhood’ (Funk, Hassan et al. 2023). We later demonstrated that shifting compartment-like voxels by 2-3 mm was sufficient to eliminate any compartment-like biases in resting state functional connectivity (Sadiq, Funk and Waugh 2025). We hypothesized that compartment-specific biases in task-based functional connectivity would also be dependent on precise voxel location. To evaluate the spatial specificity of task-evoked compartmental activation, we jittered the locations of striosome- and matrix-like voxels by ±0–3 voxels in each anatomical plane, at random and independently for each voxel. For each subject, we confirmed that although individual voxels were shifted, the mean location of the jittered masks remained nearly identical to the original compartment-specific maps. Importantly, jittered voxels did not overlap with any voxels from the original compartment-like masks. The average root-mean-square shift in individual voxel position was small: 2.9 voxels for the randomized-striosome mask and 3.1 voxels for the randomized-matrix mask. This calculation was based on the absolute magnitude of the shift. When examining the average shift within each anatomical plane (including both positive and negative displacements), the movement was minimal: 0.37 voxels on average, with a range of 0.07 to 1.2 voxels across all planes. As intended, random voxel shifts produced small but noticeable displacements at the individual voxel level yet did not meaningfully alter the overall spatial location of the full striatal masks. On average, the jittered masks still occupied the same “neighborhood” as the original striosome-like and matrix-like masks. We then used these location-shifted voxels as a negative control, comparing task-evoked activation in the original striosome- and matrix-like voxels (used in our main analyses) to that in the location-shifted voxels.

### 2.5 Analysis of task-based functional MRI (tfMRI)

We conducted a task-based fMRI analysis to examine condition-specific activation patterns within high-bias striosome-like and matrix-like striatal masks during motor task execution. First-level statistical analysis was performed using a general linear model (GLM), in which each motor task condition (hand, foot, or tongue movements) was modeled as a 12-second block, based on onset timings in the HCP-supplied explanatory variable (EV) files. Given the availability of both left-to-right (LR) and right-to-left (RL) phase-encoding acquisitions for each subject, we performed a second-level analysis to average across these runs, thereby reducing potential biases due to encoding direction. We extracted the contrast of parameter estimates (COPEs) for each task condition and computed the mean activation within striosome-like and matrix-like masks from these COPE images. Activation values represent COPE estimates derived from subject-level GLMs. These values are expressed in arbitrary units (AU), reflecting the relative amplitude of BOLD signal change associated with each condition.

In contrast to resting-state functional connectivity, which assesses spontaneous whole-brain correlation patterns, this task-based approach targeted localized, condition-locked activation and its temporal characteristics. This allowed us to directly assess how striosome- and matrix-like voxels differentially engage across the various phases of motor behavior.

### 2.6 Defining Initiation, Plateau, and Termination Phases in Motor Tasks

To assess the temporal dynamics of compartment-specific activation during movement, we segmented each motor task trial into discrete phases using the EV files provided by the HCP. These files specify the onset and duration of each 12-second movement event (e.g., hand, foot, or tongue movement) for each participant. The task includes a 3-second cue period preceding each movement block, but we did not subdivide the cue block as its short duration would not allow for sufficient temporal resolution. Instead, we focused exclusively on the 12-second movement blocks, dividing each into three equal-duration, non-overlapping, four-second windows to capture the evolving trajectory of neural activity. Specifically, the Initiation phase (0–4 s) captures activity related to movement preparation and execution; the Plateau phase (4–8 s) reflects sustained motor activity; and the Termination phase (8–12 s) represents the final portion of active movement, during which neural responses may begin to decline or transition in anticipation of rest. Notably, participants were not given a countdown or cue to signal the upcoming end of the task and were expected to continue performing movements uniformly until the end of the 12-second block. Thus, “Termination” refers to the neural dynamics associated with the tail end of the movement phase, rather than post-task deactivation. This temporal decomposition enabled us to examine whether striosome-like and matrix-like compartments exhibited distinct activation profiles across the different phases of movement execution, a key step in linking compartment-like function to behavioral timing.

### 2.7 Statistical Analysis

We assessed the accuracy of our striatal parcellations – the intra-striate position, clustering, and mean bias of our compartment-like voxels using a series of two-tailed paired-samples t-tests. We compared the volume within streamline bundles seeded by compartment-like voxels using two-tailed paired-samples t-tests.

Next, we evaluated task-induced activation levels across cue and motor conditions: cue, left foot, right foot, left hand, right hand, or tongue movement. For each task, we extracted activation values from striosome-like and matrix-like voxels separately in the left and right hemispheres. These were averaged across hemispheres to create a single mean activation value for group-level comparisons, but hemisphere-specific activation values were also retained to enable direct left–right comparisons (see Section 3.4). This resulted in two values per task per subject: one for the striosome-like compartment and one for the matrix-like compartment. For statistical comparisons of activation across movement types and cue-related activity, we conducted voxelwise analyses for each condition. To correct for multiple comparisons, we applied the Benjamini–Hochberg procedure (Benjamini and Hochberg 1995) across the family of seven tests (left foot, right foot, left hand, right hand, tongue, cue-related activity, and the average activation map across motor conditions), using a false discovery rate of Q = 0.05, resulting in a corrected significance threshold of p = 1.0×10^−5^.

For the hemispheric differences analysis, we used paired t-tests to compare activation between the left and right hemispheres for each motor task. Limb movements (hands and feet) were labeled as ipsilateral or contralateral relative to the side of movement, while tongue movement was treated as a midline task. To account for multiple comparisons across the five motor conditions, we applied the Benjamini– Hochberg false discovery rate (BH-FDR) correction with Q = 0.05, resulting in a corrected significance threshold of p = 0.014. To further quantify hemisphere-specific activation differences, we computed Laterality Indices (LI) for each motor task, defined as:

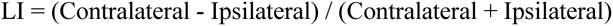

where “contralateral” refers to the hemisphere opposite the moving limb and “ipsilateral” refers to the same-side hemisphere. LI values range from −1 to +1, with positive values indicating contralateral dominance, values near zero reflecting equal bilateral or non-activation, and negative values (not observed in this study) indicating ipsilateral dominance. LI scores were computed separately for each participant and task condition, then averaged at the group level to assess task-specific lateralization trends.

Finally, we analyzed phase-specific modulation of striatal activity by dividing task-related activation into three temporal epochs: initiation, plateau, and termination. Although we computed activation values for all three phases, no significant differences between striosome-like and matrix-like voxels were observed during the initiation phase. Therefore, our primary analyses and results focus on the plateau and termination phases, where robust compartmental effects were present. For each compartment, we computed average activation within each phase and compared striosome-like and matrix-like activation levels using two-tailed paired-samples t-tests. To further characterize temporal dynamics within each compartment, we calculated linear activation slopes across phase transitions: plateau-to-termination and initiation-to-termination. We then compared these within-compartment slope values between striosome-like and matrix-like compartments using paired-samples t-tests.

## 3 Results

### 3.1 Comparing MRI-parcellated voxels to striosome and matrix in tissue

To validate our MRI-based parcellations, we first examined whether striosome-like and matrix-like voxels recapitulated their known spatial distribution within the striatum. Across diverse species from rodents to primates, the spatial distribution of striosome and matrix compartments is conserved, with striosome enriched in the rostral, ventral, and medial striatum, and matrix compartments enriched in the caudal, dorsal, and lateral striatum (Graybiel and Ragsdale Jr 1978; Goldman-Rakic 1982; Donoghue and Herkenham 1986; Ragsdale Jr and Graybiel 1990; Desban, Kemel et al. 1993; Eblen and Graybiel 1995; Waugh, Hassan et al. 2022). Our voxelwise location analysis recapitulated this compartment-specific location bias in both hemispheres. In the caudate, we found that matrix-like voxels were significantly more lateral (1.5 mm; p = 3.7×10^−51^), caudal (-8.9 mm; p < 1×10^−260^), and dorsal (8.7 mm; p < 1×10^−260^) than striosome-like voxels. In the putamen we found that matrix-like voxels were more caudal (-5.8 mm; p < 1×10^−260^) and dorsal (5.9 mm; p < 1×10^−260^) than striosome-like voxels; their medial-lateral position was not significantly different. Striosome-like voxels in the caudate were 12.2 mm medio-rostro-ventral to the centroid (Cartesian distance), while matrix-like voxels were 13.5 mm latero-caudo-dorsal to the centroid. Striosome-like voxels in the putamen were 10.2 mm medio-rostro-ventral to the centroid, while matrix-like voxels were 8.0 mm latero-caudo-dorsal to the centroid.

Histological assessments in both animal and human tissue estimated that the striosome and matrix make up approximately 15% and 85% of the striatal volume, respectively (Johnston, Gerfen et al. 1990; Desban, Kemel et al. 1993; Holt, Graybiel and Saper 1997). We evaluated compartment-like volume at a range of bias thresholds to assure that our MRI-based parcellations approximated the ratio of striosome:matrix found in tissue. Given that diffusion voxels sample the striatum in 1.25 mm cubes, independent of the underlying compartment architecture, many voxels will include both striosome and matrix tissue. Utilizing higher bias thresholds excludes more of these blended voxels and thus shifts relative volume assessments to voxels that are more striosome-like and more matrix-like. For each subject, we quantified the total number of voxels strongly biased (P > 0.87) toward striosome-like or matrix-like connectivity. Striosome-like voxels made up 5.9% of highly biased voxels, while matrix-like voxels made up 94.1%. This high cutoff excludes most striatal voxels, with 82% of the total striatal volume falling into the indeterminate category (P ≤ 0.87), reflecting an intermediate bias indicative of mixed striosome- and matrix-like connectivity. At the lowest threshold for compartment-like bias (P ≥ 0.55), striosome-like voxels constituted 25.5% and matrix-like voxels 74.5% of biased striatal volume. This lower cutoff included a greater portion of the striatum, with 45% of total striatal volume showing sufficient bias to be classified as striosome- or matrix-like, and the remaining 54% falling into the indeterminate range (P ≤ 0.55). The probability threshold that most-closely approximated the striosome-to-matrix ratio in tissue (15:85) was P ≥ 0.7, with 17.5% of voxels classified as striosome-like and 82.5% as matrix-like. This threshold included 32.2% of the total striatal volume, while the remaining 67% of voxels fell into the indeterminate category (P ≤ 0.7). These proportions varied with bias threshold but consistently reflected the marked predominance of matrix-like connectivity across the striatum. These results agree with histological evidence from both human and animal studies, demonstrating a clear preponderance of matrix-like voxels in the living human striatum.

Finally, we assessed whether striosome-like and matrix-like voxels differed in their clustering patterns. In histologic sections, each striosome branch is surrounded by contiguous matrix tissue (Graybiel and Ragsdale Jr 1978; Holt, Graybiel and Saper 1997). In contrast, our diffusion MRI voxels sampled the striatum using a rigid, non-adaptive grid that did not align with the individual-specific architecture of striosome. This misalignment introduced partial volume effects, leading to spatial blurring of striosome-like signal across adjacent voxels. Though it was not possible to resolve the fine-grained connectivity biases of individual striosome branches at this resolution, we set out to assess whether striosome-like and matrix-like voxels differed in their tendency to occur as isolated voxels or to blend into clusters. Striosome-like voxels were found in smaller, dispersed clusters, while matrix-like voxels occupied much larger and contiguous clusters. In the right hemisphere, matrix-like clusters were on average 4.3 times larger than striosome-like clusters, while in the left hemisphere, the ratio was 3.4-fold. In the right hemisphere, the mean volume of the largest striosome-like cluster was 232 mm^3^ [SEM ± 7.0; 95% CI (218, 245)], significantly smaller than the mean matrix-like cluster volume of 987 mm^3^ [SEM ± 13.2; 95% CI (962, 1013)]; p = 1.2×10⁻¹⁹⁹. Similarly, in the left hemisphere, striosome-like clusters had a mean volume of 299 mm^3^ [SEM ± 7.9; 95% CI (284, 315)], while matrix-like clusters were significantly larger, with a mean volume of 1007 mm^3^ [SEM ± 14.7; 95% CI (979, 1036)]; p = 5.6×10⁻¹⁶². These findings are consistent with histology: in tissue, the striosome is spatially dispersed and surrounded by extensive, contiguous matrix tissue.

### 3.2 Task-specific Activation Differed in Striosome- and Matrix-like Voxels

Cue-evoked and motor-evoked activation patterns showed a clear functional dissociation between striosome- and matrix-like compartments. In the cue condition, mean activation was significantly higher in striosome-like voxels (57.6 AU) than in matrix-like voxels (48.4 AU; p < 1.7×10^−11^; Figure 2). In contrast, motor-related activation was consistently higher in matrix-like voxels across all tasks. In the group-level average of all motor tasks, activation was 2.4-fold higher in matrix-like than in striosome-like voxels (matrix: 16.4, striosome: 6.9; p = 5.4×10⁻^32^). Motor-evoked activation (including both ipsilateral and contralateral hemispheres) was higher in matrix-like voxels across all movement types (Figure 2): left foot movement (matrix = 25.4, striosome = 15.1; p = 4.6×10⁻^19^); right foot movement (matrix = 22.3, striosome = 11.9; p = 1.5×10⁻^24^); left hand movement (matrix = 6.3, striosome = 2.2; p = 6.8×10⁻⁷); right hand movement (matrix = 11.6, striosome = 7.5; p = 1×10⁻⁵); tongue movement (matrix = 16.3, striosome = –2.1; p = 5.4×10⁻^32^).

**FIGURE 2:**
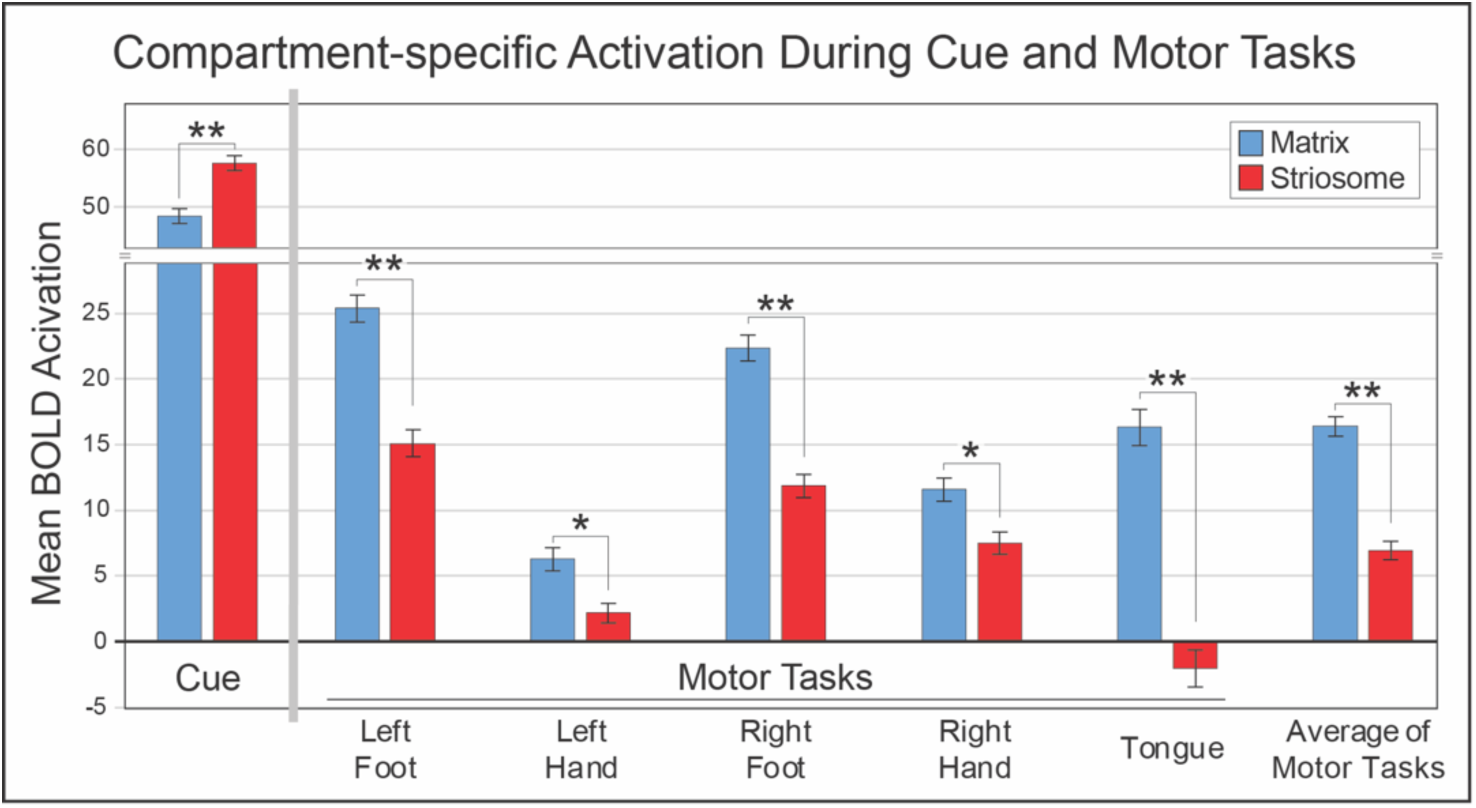
Activation differences in striosome-like and matrix-like voxels across motor tasks and cue presentation. The left-most section shows cue-evoked mean activation levels within matrix-like (blue) and striosome-like (red) compartments. Striosome-like voxels consistently had stronger cue-evoked responses. The right section shows motor-evoked mean activation levels within matrix-like and striosome-like voxels. Matrix-like voxels consistently had stronger motor-evoked responses across all tasks. Notably, tongue movement was the only task that elicited below-baseline activation in the striosome-like compartment. Error bars represent the standard error of the mean (SEM). *, p < 10^−5^. **, p < 10^−15^.

### 3.3 Influence of precise voxel location on striatal compartmentalization

Striosome and matrix occupy different parts of the striatum, on average, and so did our striosome-like and matrix-like masks. It is possible that compartment-specific differences in functional activation simply reflected the activation profiles of the parts of the striatum where striosome-like or matrix-like voxels are enriched – a “neighborhood” effect rather than a compartment-specific finding. To determine whether compartment-specific activation was dependent on the precise location of selected voxels, we compared task-based activation levels derived from the original, precisely selected striosome- and matrix-like masks (Section 3.2) to those from location-shifted versions of those compartment-like masks. We jittered the position of each compartment-like voxel individually, so the effect on the mean location of the whole mask (all voxels combined) was minimal: on average, the center of gravity of the location-shifted mask was only 0.37 voxels from the original compartment-like mask In matrix-like voxels, the average activation bias dropped from 0.95 (original) to 0.83 (shifted; paired-samples t-test, p < 1 × 10**^−260^**). In striosome-like voxels, the bias dropped from 0.78 to unbiased (0.47; paired-samples t-test, p < 1×10^−260^). These results suggest that compartment-specific activation is not explained by regional location or neighborhood effects but rather depends on the specific structural connectivity of the precisely selected voxels in our compartment-like masks. Shifting the location of striosome-like voxels reduced functional activation for cue and every motor task, significantly so in 5 of 7 task conditions (Table 1). For example, these location-shifted negative controls disrupted the expected pattern of compartmentalization: in location-shifted striosome-like voxels cue-related activation dropped by 11.8 % (p = 3.4×10^−31^) and tongue-related activation decreased by 171% (p = 4.8×10^−7^). Even though location-shifted voxels occupied the same striatal “neighborhood”, their striosome-like function was dependent on their precise location.

**Table 1:** Randomly shifting the positions of striosome-like voxels by a few voxels significantly reduced functional activation in both cue and motor tasks, while shifting matrix-like voxels did not significantly alter functional activation. Task-based activation values were compared between the original compartment-specific masks and location-shifted versions, across cue and five motor conditions. Delta activation (Shifted - Original), percentage change, and p-values reflect whether voxel displacement significantly altered mean activation within each compartment. Bolded p-values indicate significant differences (corrected for multiple comparisons). While this analysis was performed for both striosome-like and matrix-like voxels, only striosome-like voxels demonstrated significant differences in functional activation. The stability of functional activity in shifted matrix-like voxels likely reflects the abundance of matrix in the striatum; shifting the location of a matrix-like voxel is likely to select another, slightly less-biased matrix-like voxel. The pronounced changes observed in shifted striosome-like voxels highlight that compartment-specific effects depend critically on precise voxel selection rather than the “neighborhood” where a voxel is located.

| Compartment | Task | $\Delta$ Activation (Shifted – Original) | % Change | $p$-value |
| --- | --- | --- | --- | --- |
| Striosome | Cue | -6.8 | -11.8 | **$3.4 \times 10^{-31}$** |
|  | Left-Foot | -1.4 | -9.3 | **$1.0 \times 10^{-3}$** |
| | Left-Hand | -0.5 | -22.7 | $2.1 \times 10^{-1}$ |
|  | Right-Foot | -2.2 | -18.5 | **$3.0 \times 10^{-7}$** |
| | Right-Hand | -0.3 | -4.0 | $4.1 \times 10^{-1}$ |
|  | Tongue | -3.6 | -171.4 | **$4.8 \times 10^{-7}$** |
|  | Average | -1.6 | -23.2 | **$5.9 \times 10^{-7}$** |
| Matrix | Cue | 1.2 | 2.5 | $3.6 \times 10^{-2}$ |
| | Left-Foot | 0.7 | 2.8 | $9.8 \times 10^{-2}$ |
| | Left-Hand | -0.5 | -7.9 | $1.2 \times 10^{-1}$ |
| | Right-Foot | -0.7 | -3.1 | $1.0 \times 10^{-1}$ |
| | Right-Hand | -0.2 | -1.7 | $6.1 \times 10^{-1}$ |
| | Tongue | 0.3 | 1.8 | $6.2 \times 10^{-1}$ |
| | Average | -0.1 | -0.6 | $8.5 \times 10^{-1}$ |

In contrast, in the matrix-like compartment, no task condition showed a significant effect of randomization. Across tasks, Δ activation values were small (between −1.2 and +0.7) and percentage changes were consistently under 8%. This stability reflected the abundance of matrix-like voxels: shifting the location of a matrix-like voxel is likely to select another, slightly less-biased, matrix-like voxel.

Taken together, these findings provide strong evidence that striosome-like bias was spatially specific and vulnerable to randomized location shift, while matrix-like bias was stable and resistant to location shift. This negative control thus underscores the biological specificity of compartment-related differences: shifting striosome-like voxels makes them non-striosome-like, while shifting matrix-like voxels makes them only slightly less matrix-like. Compartment-specific effects depend on the precise voxel selection and are not explained by local striatal “neighborhood” activation profiles.

### 3.4 Hemispheric Differences in Striosome and Matrix Activation

We next investigated activation patterns separately in the left and right hemispheres across cue and motor tasks (Figure 3). We first combined both sets of compartment-like voxels (to identify general patterns of hemispheric asymmetry) and then assessed each set of compartment-like voxels for asymmetries. During the cue period, when activation values were averaged across both striosome-like and matrix-like voxels, the left hemisphere showed significantly greater activation (mean = 56.7) compared to the right (mean = 49.3; p < 6.8×10^−10^).

**FIGURE 3:**
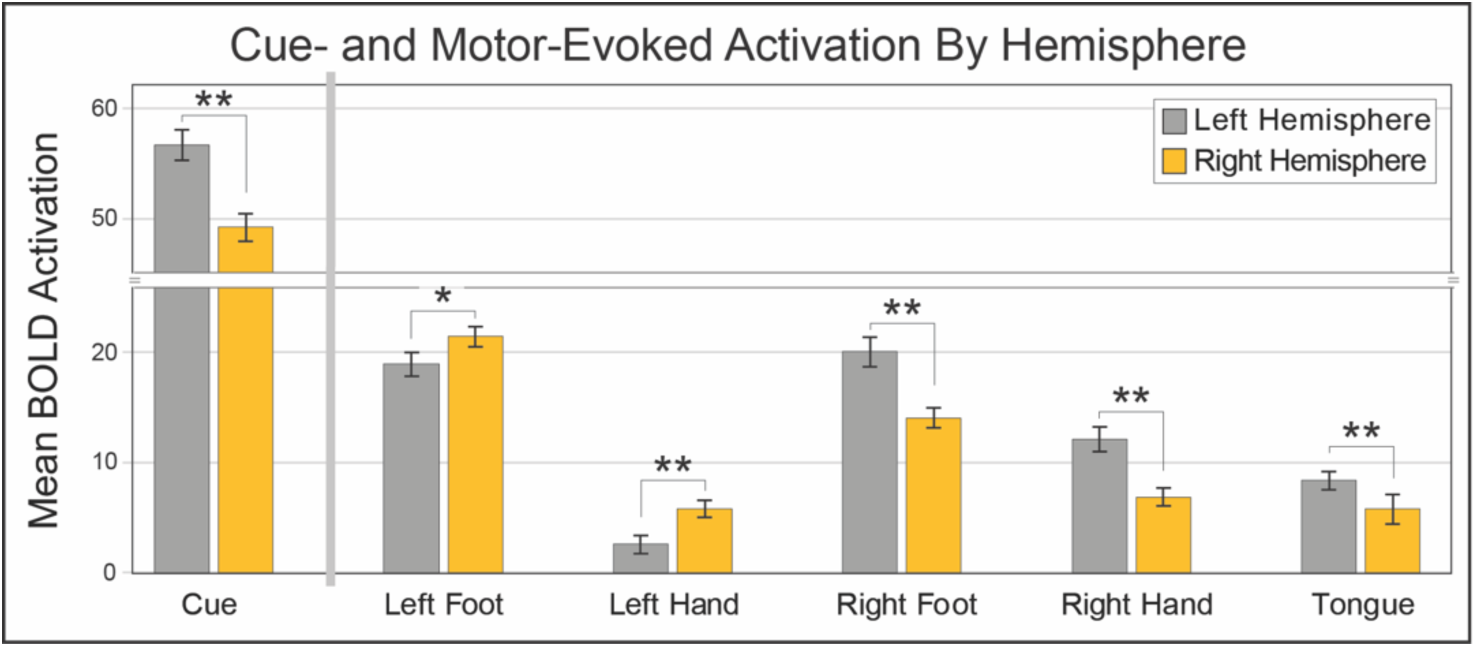
Mean activation in the left and right hemispheres during cue presentation and five motor tasks (left and right foot movement, left and right-hand movement, and tongue movement). The left side shows cue-evoked activation was significantly greater in the left hemisphere compared to the right. The right side shows that asymmetries in motor-evoked activation were evident in all motor tasks and consistently favored the contralateral hemisphere. No inter-hemispheric difference was found for tongue movements. Error bars represent the SEM. **, p < 10^−3^; *, p < 0.05.

Motor-evoked activation exhibited clear hemispheric asymmetries, particularly during hand movements (Table 2). Left- and right-hand movements preferentially activated the contralateral hemisphere. Foot movements showed a similar contralateral dominance pattern, though with slightly smaller asymmetries. In contrast, tongue movement yielded relatively symmetric activation (Table 2). In addition to contralateral dominance patterns, we observed higher overall activation magnitudes during right-sided body movements compared to left-sided ones for hand movements (total activation: right hand = 19.1, left hand = 8.4). However, the reverse was true for foot movements, where left foot activation exceeded right foot activation (total activation: left foot = 40.5, right foot = 34.2). These asymmetries may reflect a combination of motor dominance and task-specific factors such as effort or pacing.

**Table 2:** Mean activation values (including all compartment-like voxels) are shown for the left and right hemispheres during five motor tasks: left foot, right foot, left hand, right hand, or tongue movement. For limb movements, hemispheric values are labeled as either ipsilateral (Ipsi) or contralateral (Contra) relative to the body side of the movement. For tongue movement, both hemispheres are labeled as bilateral. P-values report paired t-tests comparing left vs. right hemisphere activations. Hand and foot movements exhibited strong contralateral dominance, while tongue movements evoked more symmetric responses. LH: Left Hemisphere; RH: Right Hemisphere.

| Movement Task | LH | RH | Larger Activation: | *p*-value |
| --- | --- | --- | --- | --- |
| Left Foot | 19.0 (Ipsi) | 21.5 (Contra) | Contra | 0.014 |
| Right Foot | 20.1 (Contra) | 14.1 (Ipsi) | Contra | $1.0 \times 10^{-9}$ |
| Left Hand | 2.6 (Ipsi) | 5.8 (Contra) | Contra | $1.8 \times 10^{-5}$ |
| Right Hand | 12.2 (Contra) | 6.9 (Ipsi) | Contra | $1.6 \times 10^{-10}$ |
| Tongue | 8.4 (Bilateral) | 5.8 (Bilateral) | None | 0.11 |

To quantify hemispheric asymmetries, we also computed Laterality Indices (LI) for each movement condition. Consistent with known motor system organization, hand movements showed the strongest contralateral dominance (LI = 0.38 for left hand; LI = 0.28 for right hand). Foot movements demonstrated substantially weaker lateralization overall (LI = 0.06 for left foot; LI = 0.18 for right foot), though the left–right differences were of similar numerical magnitude to those observed within hand movement trials, despite not reaching significance. Tongue movements were bilaterally represented. These patterns align with prior reports of contralateral limb representation and bilateral control of midline structures (Lotze, Montoya et al. 1999; Sörös, Schäfer and Witt 2020).

Having established overall hemispheric asymmetries, we next asked whether the degree of laterality differed between striosome-like and matrix-like voxels. Both compartments exhibited significant contralateral dominance, but this effect was stronger in matrix-like voxels. When comparing contralateral vs. ipsilateral activation across limb motor tasks (left/right hand and left/right foot), activation was significantly greater contralaterally for both striosome-like (Ipsi: 7.5, Contra: 10.8; p = 4.1×10^−8^) and matrix-like voxels (Ipsi: 13.8, Contra: 18.9; p = 8.9×10^−15^).

Contralateral activation exceeded ipsilateral activation in both compartments (Figure 4) across all motor tasks (though not significantly larger for left foot striosome). The degree of lateralization, computed as contralateral–ipsilateral difference scores, was significantly larger for matrix-like voxels than for striosome-like voxels (Striosome: 3.3, Matrix: 5.2; p = 0.03), indicating a stronger contralateral bias in matrix-like voxels.

**FIGURE 4:**
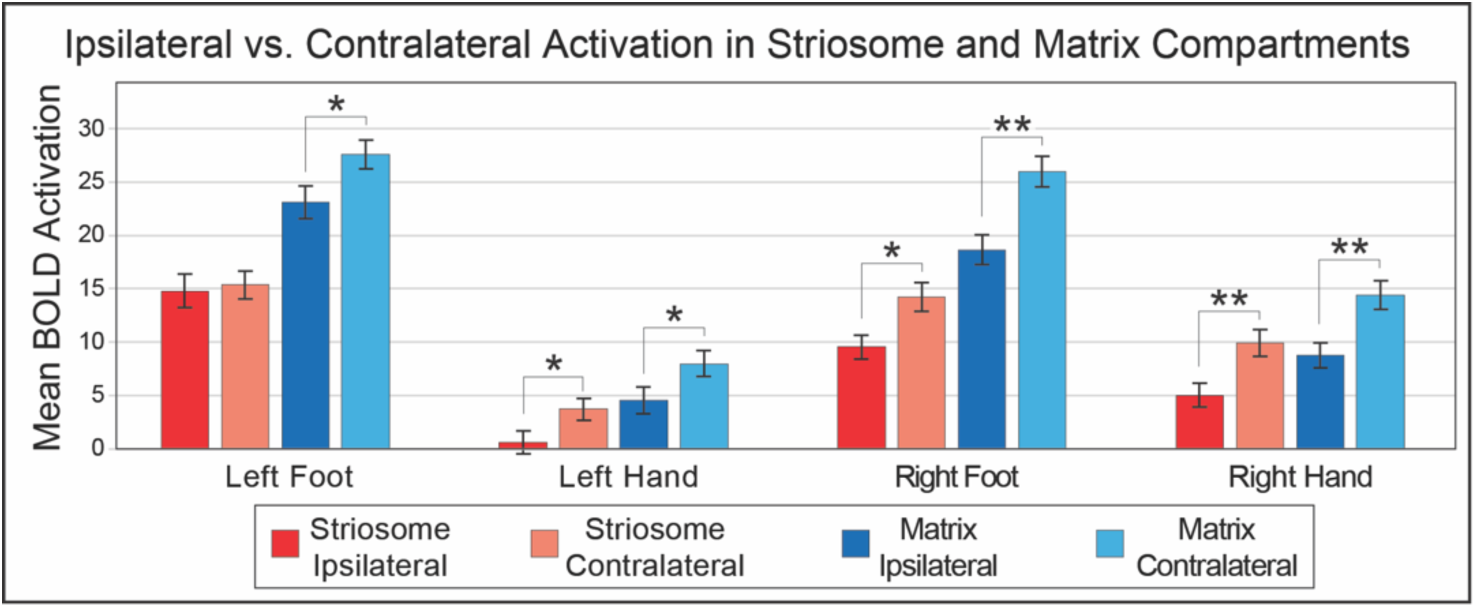
Contralateral activation exceeded ipsilateral activation in both compartments, with a stronger effect in matrix-like voxels. Ipsilateral vs. contralateral activation in striosome-like and matrix-like voxels during motor tasks. Bars represent mean activation (BOLD signal change) across all tasks for each compartment and hemisphere condition. Error bars indicate SEM. *, p < 10^−3^; **, p < 2×10^−5^.

### 3.5 Phase-Specific Modulation of Striosome- and Matrix-like Activation

We found different patterns of phase-specific activation between striosome-like and matrix-like voxels, particularly during the transition from plateau to termination (Figure 5A). These effects reflect dynamic compartment-specific modulation of activity during motor execution. During the initiation phase, both compartments exhibited similarly high activation (striosome: 29.7, matrix: 30.1), which may reflect equivalent recruitment at the onset of movement or residual activation from the preceding cue block, which was substantially higher than activation in the motor tasks (Fig. 2). In the plateau phase, activation declined moderately in both compartments (striosome-like: 14.1, matrix-like: 15.4), reflecting waning functional activation. However, during the termination phase, activation trajectories diverged: activation in striosome-like voxels dropped below baseline (–1.7), while activation in matrix-like voxels remained positive (9.3; p < 1.3×10^− 10^), indicating more sustained involvement of matrix-like voxels at task offset.

**FIGURE 5:**
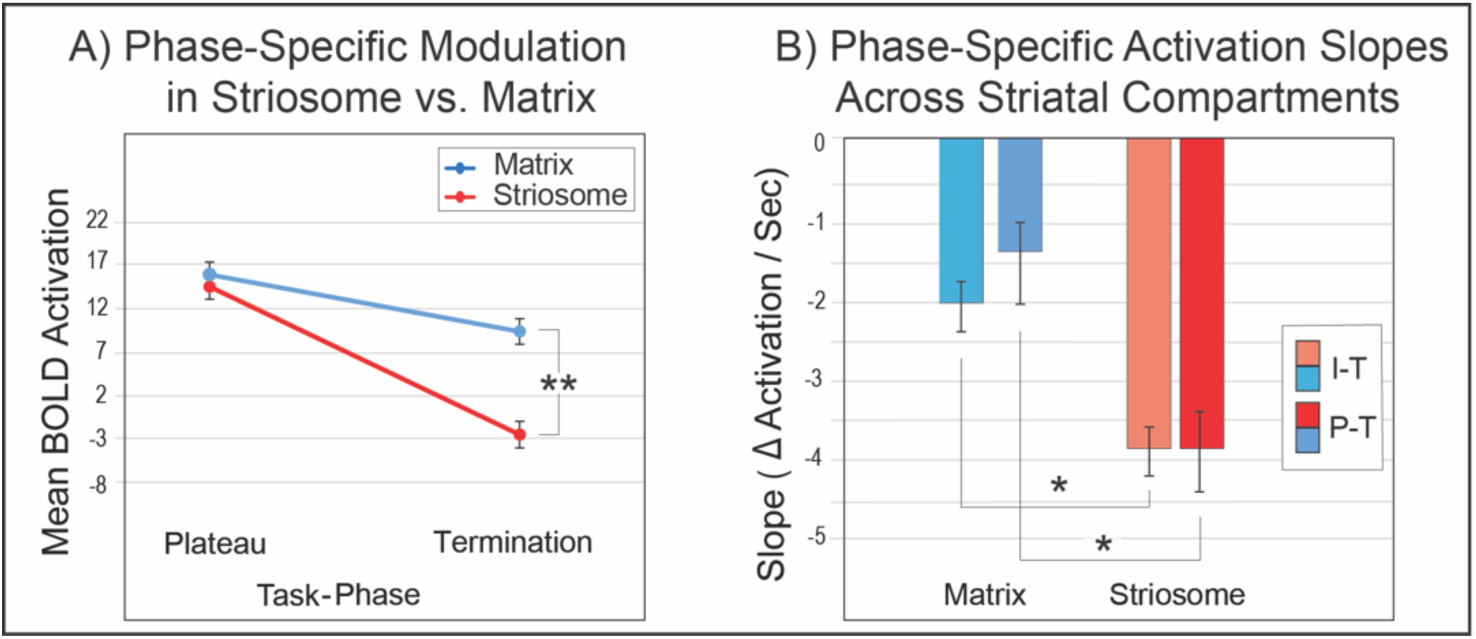
Matrix-like voxels had more sustained activation during motor execution compared to striosome-like voxels. We divided each 12 second movement block into three phases: initiation, plateau, and termination. The initiation phase was heavily influenced by residual activation from the high-activation cue block (Fig. 2), leading us to focus on the plateau and termination phases of movement. (A) Mean activation in the plateau and termination phases for striosome-like and matrix-like voxels. (B) Activation slopes between task phases (plateau to termination [P-T], and initiation to termination [I-T]) illustrating dynamic changes in compartment-like activity over time. Error bars represent SEM. **, p=1.3×10^−10^; *, p < 7×10^−4^.

To quantify temporal differences in activation patterns, we computed phase-specific slopes for each compartment-like mask, representing the rate of activation change across task phases (Figure 5). From initiation to plateau, striosome-like and matrix-like voxels exhibited similar declines in activation (striosome slope: –3.9; matrix slope: –3.7), and the difference between them was not significant (p = 0.7). However, from plateau to termination, the slopes diverged sharply: activation in striosome-like voxels showed a steep deactivation (–4.0), while in matrix-like voxels, activation declined more gradually (–1.5; *p* = 1.3×10⁻^4^). This difference in the slope of deactivation was also evident when comparing the first to the last phase (initiation to termination): striosome-like voxels again exhibited a steeper overall decline in activation (–3.9) than matrix-like voxels (– 2.6; p = 6.5×10^−4^). These results highlight a biphasic divergence in temporal dynamics: matrix activation decreased gradually across phases, whereas striosomal activation dropped more abruptly at task termination.

## 4 Discussion

The fundamental differences in the embryology, pharmacology, and connectivity of the striosome and matrix suggest that their functions in behavior may also be divergent (Graybiel and Ragsdale Jr 1978; Brimblecombe and Cragg 2017; Prager, Dorman et al. 2020). Indeed, distinct, compartment-specific functional roles have been demonstrated in an array of animal behaviors: reward (White and Hiroi 1998), decision making under threat (Friedman, Homma et al. 2015; Amemori, Graybiel and Amemori 2021), habit formation (Nadel, Pawelko et al. 2021), learning (Jenrette, Logue and Horner 2019;

Xiao, Deng et al. 2020), and motor control (Weglage, Wärnberg et al. 2021; Okunomiya, Watanabe et al. 2025). However, to the best of our knowledge, distinct functional roles for striosome and matrix have never been identified in living humans.

Our findings demonstrate that the striosome and matrix compartments exhibit temporally distinct activation patterns during motor behaviors. Striosome-like voxels were preferentially active during motor preparation, whereas matrix-like voxels showed stronger activation during motor execution. This temporal dissociation mirrors compartmental roles inferred from animal studies, in which striosomal circuits are preferentially engaged during decision-making, reward anticipation, and behavioral inhibition, while matrix circuits are more active during motor initiation and execution (Graybiel 2008; Hikosaka, Kim et al. 2014; Friedman, Homma et al. 2015; Okunomiya, Watanabe et al. 2025) and suggests that the human striatum maintains functional segregation consistent with its microanatomical organization. These results build on our prior resting-state work (Sadiq, Funk and Waugh 2025), which showed that striosome and matrix occupy distinct intrinsic functional networks; here we show that they also differ in task-evoked dynamics. Activation in striosome-like voxels peaks during preparation, whereas matrix-like voxels peak during motor execution. The fact that this dual organization is present at rest and during behavior implies that compartment-based functional specialization is a core feature of striatal architecture. Finally, our data raises the possibility that imbalances in compartment-specific engagement may contribute to motor symptoms observed in neuropsychiatric disorders (Tippett, Waldvogel et al. 2007; Crittenden and Graybiel 2011; Waugh, Hassan et al. 2025).

It is important to acknowledge several limitations of the study. First, our approach identified striosome- and matrix-like voxels indirectly, using biases in structural connectivity identified through probabilistic tractography. The accuracy of our functional assessments depends on the anatomical accuracy of our striatal parcellations. We previously demonstrated that striatal parcellation is highly reliable (test-retest error rate of 0.14% (Waugh, Hassan et al. 2022) and that these biased patterns of connectivity depend on highly precise selection of striatal voxels – shifting voxel position by just a few millimeters negates all compartment-like bias (Funk, Hassan and Waugh 2024; Sadiq, Funk and Waugh 2025). Moreover, striatal parcellation recapitulates all of the anatomical features of striosome and matrix demonstrated in human and animal tissue: their relative abundance, spatial distribution, contiguity, and extra-striate connectivity (Waugh, Hassan et al. 2022; Funk, Hassan et al. 2023; Funk, Hassan and Waugh 2024). However, this technique is not the equivalent of direct identification of the compartments through histological staining; compartment-like voxels share many features with the striosome and matrix, but the degree to which connectivity-based parcellation approximates the compartments is unknown.

A further limitation is the spatial resolution of diffusion MRI. Even though the voxel resolution used in this study (1.25 mm isotropic) matches the diameter of the largest human striosome branches (Graybiel and Ragsdale Jr 1978; Holt, Graybiel and Saper 1997), every striosome-like voxel will include some fraction of matrix tissue. However, it is notable that our previously-described validation experiments – in which compartment-like voxels matched the anatomic features of striosome and matrix identified through immunohistochemistry – were accurate in diffusion datasets with lower resolution than we utilized here (Waugh, Hassan et al. 2022; Funk, Hassan et al. 2023; Funk, Hassan and Waugh 2024). The resolution of our fMRI dataset matches that of our prior diffusion datasets (2 mm isotropic), and we demonstrated that this resolution is sufficient to identify widespread and robust compartment-specific patterns of functional connectivity (Sadiq, Funk and Waugh 2025). However, we urge readers to recall that these inferential methods cannot match the fine-grained histological detail available in post-mortem tissue. This limitation is inherent to all current non-invasive anatomical mapping techniques.

The unique distribution of the striosome within each person’s striatum precludes the use of region-of-interest based assessments. Therefore, we analyzed mean activation within the striosome-like and matrix-like masks for each subject, rather than conducting voxel-wise activation analyses. While averaging within each compartment enabled meaningful between-subjects comparisons, this approach may have obscured more spatially localized activation patterns within each compartment. Since corticostriate projections are organized somatotopically (Flaherty and Graybiel 1993; Waugh, Hassan et al. 2022; Sadiq, Funk and Waugh 2025), the functional specializations we identified may be localized to particular parts of each striatal compartment.

It is also important to consider ways in which our experimental approaches may have distorted functional connectivity. Histological studies consistently report that matrix occupies approximately 6-fold larger volume than striosome (Desban, Kemel et al. 1993; Holt, Graybiel and Saper 1997). However, our goal of generating equal-volume masks, which was necessary to avoid size-based biases in probabilistic connectivity, also led us to assess only a fraction of all matrix-like voxels. It is possible that other matrix-like voxels, those not among the most-biased set, could have different functional activation patterns than the ones we identified. The natural volume asymmetry between striosome and matrix remains relevant for understanding the anatomical context of our findings but does not diminish the functional activation patterns reported here.

Our task-based fMRI results revealed a clear functional dissociation between striosome-like and matrix-like voxels. During the cue period, striosome-like voxels exhibited significantly higher activation than matrix-like voxels, suggesting a specialized role in reading, directing attention, or anticipatory processes such as motor preparation, decision-making, or motivational evaluation, functions that have been previously attributed to the striosome (Friedman, Homma et al. 2017; Karunakaran, Amemori et al. 2021). In contrast, motor execution elicited greater activation in matrix-like voxels across all movement types, providing the first direct functional evidence in humans for the matrix’s proposed involvement in motor execution, a role that has been inferred primarily from cortico-striate structural connectivity (Flaherty and Graybiel 1993). Notably, task-evoked activation amplitudes in matrix-like regions varied systematically by effector: activation was strongest during foot movements, followed by tongue, and lowest during hand movements. This pattern may reflect differences in how demanding or effortful these movements are, the degree to which a movement engages with postural or positional stabilizing muscles, or may be proportionate to the volume of muscle activated to achieve these movements. These results suggest that matrix-favoring regions may be more strongly engaged for movements that involve greater motor drive or larger fractions of the motor homunculus. Overall, our findings support the idea that striosome and matrix compartments play distinct functional roles in movement tasks depending not only on the phase of the task (cue vs. movement) but also on which body part is being moved.

Interestingly, the reduced or even negative activation of the striosome during overt motor tasks, most notably during tongue movement (Fig. 2), underscores a possible compartmental specialization within tasks, with different functions during preparatory and execution phases. One possible explanation is that striosome activity supports early-phase processes such as movement initiation, anticipatory postural adjustment, or context evaluation, and gradually disengages as the movement unfolds, particularly during the termination phase. This temporal pattern suggests that striosomal circuits may contribute most during the transition into action and the regulation of when to stop or adjust behavior, rather than during sustained execution. The observed pattern of striosomal activation (strongest for foot movements, moderate for hand movements, and lowest for tongue movements) may reflect how routine or automatic each type of movement is. Foot movements, particularly when not part of walking or other reflex-triggered actions, may require more mental planning. In contrast, hand movements are often well-practiced and automatic. Tongue movements in this context may be the simplest and most repetitive, requiring the least preparation. Another possibility is that striosome activation correlates with the need for postural stabilization or adjustment, with foot greater than hand, and tongue requiring no postural adjustment. These findings align with and extend prior animal research that suggested that striosome activity in humans may be more closely tied to motor planning, postural preparation, and context-dependent modulation (Graybiel and Matsushima 2023), while matrix activity may support the execution of well-learned or habitual motor actions (Redgrave, Rodriguez et al. 2010). The observed activation profiles further imply that disruptions in inter-compartmental balance could contribute to the motor, limbic, or cognitive symptoms seen in disorders such as Parkinson and Huntington diseases, where striosome and matrix are differentially impacted (Tippett, Waldvogel et al. 2007; Crittenden and Graybiel 2011; Marecek, Krajca et al. 2024). Notably, while our prior resting-state fMRI analysis revealed stronger intrinsic connectivity in striosome-like voxels (Sadiq, Funk and Waugh 2025), the current task-based findings show that matrix-like voxels are more strongly engaged during motor execution. This pattern reinforces the idea that striosome and matrix compartments contribute differentially to intrinsic (resting) and evoked (task-based) brain states, consistent with complementary roles in selecting and preparing for actions versus action implementation.

In addition to overall hemispheric asymmetries (Figure 3), we also examined whether these effects differed between striosome- and matrix-like voxels (Figure 4). When averaged across compartments, cue-related activation was stronger in the left hemisphere, potentially a correlate of prior evidence linking the left striatum to internal processing and self-control (Zhang, Hu et al. 2017). In contrast, during motor execution, matrix-like voxels showed task-specific contralateral dominance, with greater right than left hemisphere activation.

When hemispheric effects were analyzed separately by compartment (Figure 4), both striosome- and matrix-like voxels exhibited significant contralateral > ipsilateral activation. However, the effect was significantly stronger in matrix-like voxels, which showed higher absolute activation levels during all motor tasks. These task-based lateralization patterns align with our prior resting-state findings, where the right striatum displayed stronger compartment-specific functional connectivity biases (Sadiq, Funk and Waugh 2025). Notably, our interpretation is supported by tract-tracing work from Fujiyama et al. (Fujiyama, Unzai and Karube 2019), who assessed bilateral corticostriatal projections from motor cortical areas in non-human primates. They found that proximal limb and postural areas give rise to substantial bilateral input to the striatum, suggesting an anatomical substrate for interhemispheric coordination. Together, these converging findings imply that striosome and matrix contribute to lateralized and possibly bilaterally integrated control of motor behavior, with the striosome playing a greater role in preparatory and internally driven processes and the matrix in contralateral motor execution and coordination.

To further characterize compartment-specific specialization, we computed differential activation indices across tasks. This analysis revealed a clear division of labor: striosome compartments were more active during the cue period, while matrix compartments consistently dominated during motor execution, with the strongest preference observed for tongue movements, followed by foot and hand tasks. These results align with prior evidence that striosomal neurons encode predictive information early in trials (Yoshizawa, Ito and Doya 2018). For instance, Yoshizawa et al. showed in mice that increased striosomal activity rose in response to reward-predictive cues, and that activation was proportional to expected reward value and persisted after controlling for motor outputs, such as licking. Similarly, Bloem et al. (2017) found a higher proportion of striosomal neurons responded selectively to reward-predicting auditory tones, with this preference strengthening over the course of training (Bloem, Huda et al. 2017). These findings highlight the striosome’s specialized role in anticipatory processing, and our task-based fMRI data provide the first evidence that this cue-period specialization extends to humans.

Further supporting the compartmental distinction, Bloem et al. (2022) demonstrated that striosomal neurons preferentially encoded reward and punishment prediction errors in a cost–benefit decision-making task. Even in the absence of explicit external cues, striosomal neurons were preferentially active during early anticipatory phases, when animals relied on internal models to predict outcomes (Bloem, Huda et al. 2022). This finding reinforces the role of the striosome in encoding internal evaluations during the preparatory stages of action selection. Additionally, Kim et al. (2024) reported sustained dopamine release during the cue period of a discrimination task, with plateau-like dynamics in centromedial striatal regions, an area where the striosome is abundant (Kim, Gibson et al. 2024). While their study did not directly distinguish compartments, the observed dopamine dynamics are consistent with the notion that cue-period dopaminergic signaling supports anticipatory processing, aligning with our observation of striosomal activation during cues.

Our findings revealed distinct phase-specific activation profiles between striosome- and matrix-like voxels during motor task execution. Matrix-like voxels showed robust activation during the initiation phase, which declined only modestly through the plateau and termination phases. These findings align with prior work suggesting the dorsolateral striatum (DLS) plays a key role in the execution and initiation of habitual sequences (Jin and Costa 2010). Although Jin and Costa did not examine striosome and matrix compartments separately, the DLS is known to be matrix-dominant (Graybiel and Ragsdale Jr 1978; Goldman-Rakic 1982; Donoghue and Herkenham 1986; Gimenez-Amaya and Graybiel 1990; Desban, Kemel et al. 1993; Eblen and Graybiel 1995), suggesting that the observed role of the DLS in habitual behavior may be mediated largely by matrix circuits. As skills are learned and become automatic, striatal involvement shifts from associative (dorsomedial striatum) to sensorimotor (DLS) circuits, a transition supported by region-specific synaptic plasticity, particularly in the indirect pathway of the DLS. In contrast striosome-like voxels showed comparable activation to matrix-like voxels at the plateau but exhibited a pronounced drop during the termination phase. This pattern suggests that striosome engagement is maintained through initiation and plateau, with a selective reduction as the movement concludes. This temporal divergence was further quantified in slope analyses: striosome activation remained stable between initiation and plateau but dropped significantly from plateau to termination while matrix activation declined steadily across all phases (Fig. 5).

Another possibility is that the pronounced drop in striosomal activation during task termination may reflect a functional disengagement once evaluative or predictive demands subside, potentially signaling behavioral transitions or the resolution of action plans. This interpretation aligns with anatomical findings that striosomal neurons receive input from limbic regions involved in internal state monitoring and project to midbrain dopaminergic areas implicated in behavioral regulation and state transitions (Fuccillo 2016; Brimblecombe and Cragg 2017). Supporting a role in action modulation, Okunomiya et al. (2025) showed that prolonged chemogenetic activation of striosomal neurons via DREADDs led to sustained motor slowing in mice, suggesting that elevated striosome activity may inhibit movement execution (Okunomiya, Watanabe et al. 2025). Notably, their manipulation spanned minutes, far exceeding the transient 4-second termination window in our task. Our findings suggest a temporally confined role for striosomal disengagement during transitions rather than generalized inhibition. This divergence in slopes supports a compartmental division of labor: matrix circuits contribute to sustained execution, while striosomal circuits regulate transitions, boundary marking, and internal evaluation, possibly through their direct inhibition of nigral dopaminergic neurons (Crittenden, Tillberg et al. 2016).

Importantly, we also observed hemispheric asymmetries in both compartments. Striosome activation during cue phases was stronger in the left hemisphere, consistent with the left striatum’s proposed role in internal monitoring and self-regulation (Zhang, Hu et al. 2017). The stronger left-hemisphere activation during the cue phase may partly reflect language-related processing, such as reading the cue, given the typical left-hemisphere dominance for language (Tzourio-Mazoyer, Josse et al. 2004; Price 2012). In contrast, matrix activation during motor execution exhibited task-specific contralateral dominance, particularly for foot and hand movements, mirroring classical patterns of lateralized motor control (Draganski, Kherif et al. 2008). Together, these findings reinforce the idea that the striosome and matrix compartments are not only functionally segregated in time but also organized across hemispheres to support preparatory versus execution roles in motor behavior.

In conclusion, this study provides the first task-based fMRI evidence that compartment-like voxels in the human striatum exhibit distinct, temporally patterned activation profiles during motor behaviors. While our prior resting-state fMRI work demonstrated that these compartment-like voxels participate in segregated intrinsic functional networks (Sadiq, Funk and Waugh 2025), the current findings suggest that such compartmental organization also influences activation dynamics during task performance. These results support the emerging view that striatal microarchitecture may contribute to phase-specific functional specialization, with striosome- and matrix-like voxels differentially engaged across cue processing and motor execution. Further studies will be needed to directly map these functional signatures to underlying cellular and circuit-level mechanisms, and to explore their relevance across cognitive domains and clinical populations.

By integrating structural connectivity-based parcellation, temporal decomposition of task epochs, and activation profiling, our findings extend the functional characterization of the striatal compartments beyond resting-state connectivity to real-time functional activation during motor behaviors. This framework enhances our understanding of how striosome- and matrix-specific networks support different facets of human behavior. These findings also suggest that compartment-specific maldevelopment or injury may explain distinct motor features of neuropsychiatric disorders that involve the striatum or striatal networks.

## Data availability statement

Publicly available datasets were analyzed in this study. HCP data can be found here: https://www.humanconnectome.org/study/hcp-young-adult/document/1200-subjects-data-release. This dataset is BIDS compliant. The code, bait, seed, and exclusion masks necessary to complete striatal parcellation can be accessed here: github.com/jeff-waugh/Striatal-Connectivity-based-Parcellation.

## Author Contributions

AS: data acquisition and analysis, initial manuscript drafting, and critical revision of the manuscript. JW: data acquisition, analysis, and interpretation, initial manuscript drafting, and critical manuscript revision. All authors contributed to the article and approved the final version for submission.

